# ALPHLARD: a Bayesian method for analyzing HLA genes from whole genome sequence data

**DOI:** 10.1101/323766

**Authors:** Shuto Hayashi, Rui Yamaguchi, Shinichi Mizuno, Mitsuhiro Komura, Satoru Miyano, Hidewaki Nakagawa, Seiya Imoto

## Abstract

Although human leukocyte antigen (HLA) genotyping based on amplicon, whole exome sequence (WES), and RNA sequence data has been achieved in recent years, accurate genotyping from whole genome sequence (WGS) data remains a challenge due to the low depth. Furthermore, there is no method to identify the sequences of unknown HLA types not registered in HLA databases. We developed a Bayesian model, called ALPHLARD, that collects reads potentially generated from HLA genes and accurately determines a pair of HLA types for each of HLA-A, -B, -C, -DPA1, -DPB1, -DQA1, -DQB1, and -DRB1 genes at 6-digit resolution. Furthermore, ALPHLARD can detect rare germline variants not stored in HLA databases and call somatic mutations from paired normal and tumor sequence data. We illustrate the capability of ALPHLARD using 253 WES data and 25 WGS data from Illumina platforms. By comparing the results of HLA genotyping from SBT and amplicon sequencing methods, ALPHLARD achieved 98.8% for WES data and 98.5% for WGS data at 4-digit resolution. We also detected three somatic point mutations and one case of loss of heterozygosity in the HLA genes from the WGS data. ALPHLARD showed good performance for HLA genotyping even from low-coverage data. It also has a potential to detect rare germline variants and somatic mutations in HLA genes. It would help to fill in the current gaps in HLA reference databases and unveil the immunological significance of somatic mutations identified in HLA genes.

## Introduction

Human leukocyte antigen (HLA) genes play a key role in immunological responses by presenting peptides to T cells. It is well known that HLA loci are highly polymorphic, and the polymorphism patterns define several thousands of types within HLA genes. HLA genotyping is a process that determines a pair of HLA types for an HLA gene. Since the relationships between HLA types and diseases have now been intensively investigated [1–5], HLA genotyping is considered as a fundamental step in immunological analysis. Further analysis enables us to identify novel HLA types and detect somatic mutations, which potentially affect the efficacy of immune therapy.

Recently, next generation sequencing-based approaches have been developed for HLA genotyping. These can be generally separated into two categories: those based on amplicon sequencing of HLA loci [6, 7] and others based on unbiased sequencing methods such as whole exome sequencing (WES) and RNA sequencing (RNA-seq) [8–15]. The amplicon sequencing-based methods are the most accurate owing to the sufficient coverage of sequence data, but are relatively expensive to perform and require specialized materials and equipment. The unbiased sequencing ones can be used without additional costs, but the accuracy of the results depends on the amount and quality of sequence reads generated from HLA loci. Previous papers have shown that the accuracy can reach 95% at 4-digit resolution from WES and RNA-seq data [10, 12, 13, 15] However, Bauer *et al.* has reported that these methods cannot achieve 80% accuracy from whole genome sequence (WGS) data [16]. Thus, HLA genotyping from WGS data remains a significant challenge, although this approach would provide more information of HLA loci than possible with WES and RNA-seq data, including details of the non-coding regions such as the introns and the untranslated regions.

To achieve high accuracy for WGS-based HLA genotyping and further analysis of HLA genes, we developed a series of computational methods, which involve collection of sequence reads that are potentially generated from a target HLA gene followed by HLA genotyping, using a novel Bayesian model termed ALelle Prediction in HLA Regions from sequence Data (ALPHLARD). This model was found to yield comparable accuracy to those based on WES and RNA-seq data at 6-digit resolution. Together with HLA genotyping, a notable feature of ALPHLARD is that it can estimate the personal HLA sequences of the sample. This enables achieving high accuracy for a sample whose HLA sequence is not included in the reference databases and further allows for calling rare germline variants not stored in the databases. We can also detect somatic mutations by comparing the HLA sequences of paired normal and tumor sequence data.

We illustrate the capability of our method by comparing the performance of ALPHLARD and existing methods using WES data from 253 HapMap samples and WGS data from the normal samples of 25 cancer patients. We also applied ALPHLARD to WGS data of the tumor samples of the cancer patients and detected three somatic point mutations and one case of loss of heterozygosity (LOH) in the HLA genes, which were validated by the Trusight HLA Sequencing Panels [17] and the Sanger sequencing.

## Methods

### Overview of our pipeline

Our pipeline consists of two steps as shown in Figure 1. First, for each read and each HLA type, the HLA read score (HR score) is calculated, which quantifies the likelihood that the read comes from the HLA type. Based on the calculated HR scores, it is determined whether or not the read comes from a certain HLA gene. For example, by aligning read *x* to the reference sequences in HLA databases, we obtained the HR scores as shown in the bar graph of Figure 1a. Then, if the maximum HR score for the HLA-A gene is large enough and the difference in the maximum scores for the HLA-A gene and the other HLA genes is also large, we conclude that read *x* is most likely a specific read of the HLA-A gene. Otherwise, read *x* is judged to be a read produced from other regions. HLA genotyping is then performed using the collected reads for each HLA gene, as shown in Figure 1b. ALPHLARD outputs candidate pairs of HLA types according to the Bayesian posterior probabilities.

**Figure 1:**
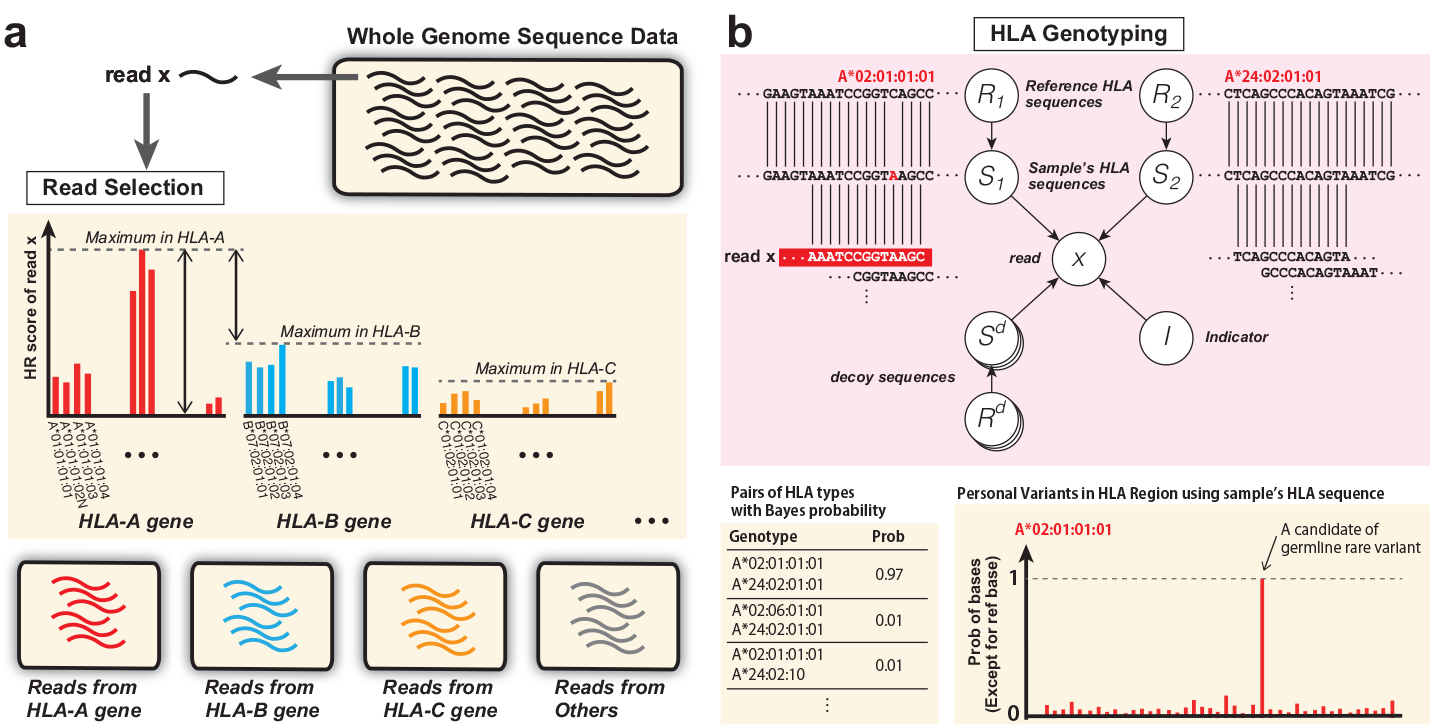
Schematic overview of ALPHLARD: (a) For each read and each HLA type, the HLA read score (HR score) is calculated, which quantifies the likelihood that the read comes from the HLA type. Based on the calculated HR scores, it is determined whether or not the read comes from a certain HLA gene. (b) For each read and each HLA type, the HLA read score (HR score) is calculated, which quantifies the likelihood that the read comes from the HLA type. Based on the calculated HR scores, it is determined whether or not the read comes from a certain HLA gene.

### HLA reference data

We used HLA reference information that can be obtained from the IPD-IMGT/HLA database (release 3.28.0) [18]. There are two types of HLA reference sequences in the database: one is a complete genomic reference and the other is an exonic reference without non-coding regions. Some HLA types have both genomic and exonic reference information, but most HLA types have only exonic reference information.

The database also provides multiple sequence alignments (MSAs) at the genomic and the exonic levels for each HLA gene. We combined the two MSAs into a common MSA as follows: First, some gaps were inserted into exons of the genomic MSA for consistency with the exonic reference sequences. Then, missing non-coding sequences were replaced with the most similar genomic reference sequences. This integrated MSA is then used for alignment and realignment of the reads.

### Collection and realignment of reads

First, all reads are mapped to a human reference genome, and reads mapped to the HLA region and unmapped reads are used at the next step. We use hg19 [19] as the reference sequence and define the HLA region as chr6:28,477,797-33,448,354, which covers HLA-A, -B, -C, -DPA1, -DPB1, -DQA1, -DQB1, and -DRB1 genes.

Next, the filtered reads are mapped to all HLA genomic and exonic reference sequences. We use BWA-MEM (version 0.7.10) [20] with the -a option to output all found alignments. Then, each mapped read is filtered based on whether or not it is likely to be produced by the target HLA gene. This filtering is performed according to the HR score *s*_*ij*_ for the *i*^*th*^ read *x*_*i*_ and the *j*^*th*^ HLA type *t*_*j*_, which is similar to the filtering procedure used in HLAforest [11]. If *x*_*i*_ is not aligned to *t*_*j*_, *s*_*ij*_ is —∞. Otherwise, let (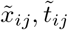) be the alignment of *x*_*i*_ and *t*_*j*_, which might include some gaps. 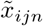 and 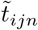 are defined as the *n*^*th*^ bases or gaps of 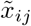 and 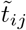, respectively, and 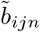 is defined as the base quality of 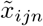. We suppose that 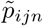 is the probability of a mismatch between 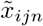 and 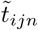, which can be calculated by

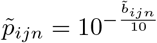

Then, the HR score *s*_*ij*_ is given by where

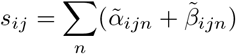

Where

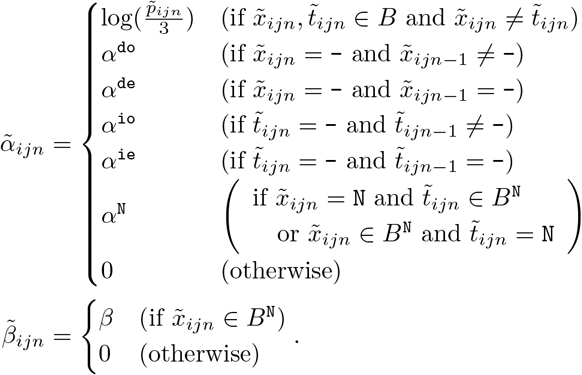

Here, *B* = {A, C, G, T} and *B*^N^ = {A, C, G, T, N}. The parameters *α*^do^, *α*^de^, *α*^io^, *α*^ie^, and *α*^N^ take negative values as penalties for opening a deletion, extending a deletion, opening an insertion, extending an insertion, and N in the read or the HLA type, respectively. *β* is a positive constant reward for read length, which prefers longer reads. Then, the score of *x*_*i*_ for the target HLA gene 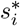, and the score of *x*_*i*_ for the non-target HLA genes 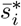 are defined by

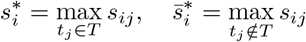

where *T* is the set of HLA types in the target HLA gene. 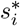 and 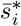 indicate how likely *x*_*i*_ is to be produced by the target HLA gene and the non-target HLA genes, respectively.

Thus, when *x*_*i*_ is an unpaired read, it is used for HLA genotyping if

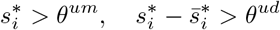

where *θ*^*um*^ and *θ*^*ud*^ are constant thresholds. When *x*_*i*_ and *x*_*i*′_ are paired, they are used for HLA genotyping if

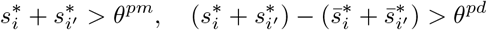

where *θ*^*pm*^ and *θ*^*pd*^ are constant thresholds. Paired reads are generally more effective than unpaired reads; hence, *θ*^*pm*^ and *θ*^*pd*^ should be less than *θ*^*um*^ and *θ*^*ud*^, respectively.

In the next step, all of the collected reads are realigned as follows. First, *t*_*j*\*_ is defined as the best type for *x*_*i*_ in the target gene, which is obtained by

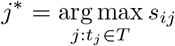

Then, *x*_*i*_ is realigned to be consistent with the alignment (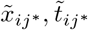) and the integrated MSA of the target HLA gene.

### Bayesian model for analyzing HLA genes

Analysis of the target HLA gene by ALPHLARD is performed using the collected and realigned reads. Let 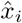 be the *i*^*th*^ paired (or unpaired) read(s) collected and realigned with the previous procedure, 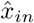 be the *n*^*th*^ base or gap of 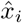, and 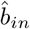 be the base quality of 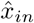. Note that, hereafter, we regard paired reads as one sequence. The probability of mismatch 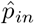 can be calculated by

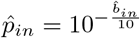

Suppose that 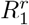 and 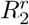 are the HLA types of the sample, and that 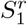 and 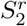 are the true HLA sequences of the sample, which are introduced because the HLA sequences of the sample might not be registered in the reference (IPD-IMGT/HLA) database. Let 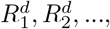 be decoy HLA types and 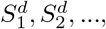 be decoy HLA sequences. These parameters could make this HLA analysis robust when reads from non-target homologous regions are misclassified into the target HLA gene at the previous filtering step. We will sometimes use *R*_1_,*R*_2_,*R*_3_,*R*_4_,…, and *S*_1_,*S*_2_,*S*_3_,*S*_4_,…, instead of 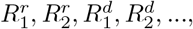 and 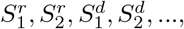 for convenience. *I*_*i*_ is defined as a parameter to indicate which sequence produced the read 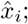 that is, *I*_*i*_ = *k* means that 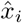 was generated from *S*_*k*_. Then, the posterior probability of the parameters 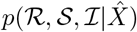 is given by

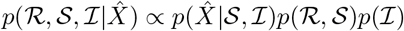

where 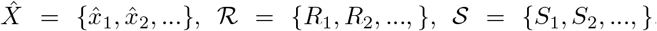, and 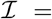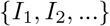.
The likelihood function 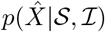 is defined by

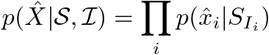

The likelihood of each read 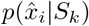 is given by

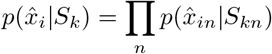

where *S*_*kn*_ is the *n*^*th*^ base or gap of *S*_*k*_. The likelihood of each base 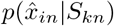
is calculated by

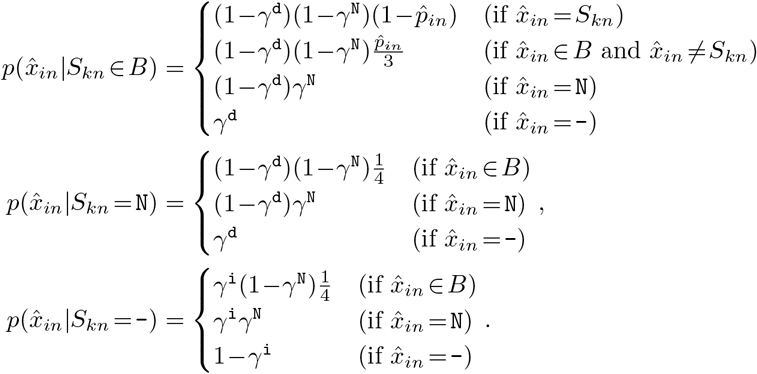

Here, γ^d^, γ^i^, and γ^N^ are the probabilities of a deletion error, an insertion error, and N, respectively.

The prior probability of the HLA types and the HLA sequences 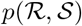 is defined by

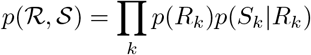

Here, 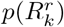 is the prior probability of the HLA type, which is calculated using The Allele Frequency Net Database [21]. On the other hand, 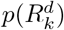 is the prior probability of the decoy HLA type, which we assume as constant. The prior probability of the HLA sequence *p*(*S*_*k*_|*R*_*k*_) is given by

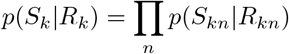

where and *R*_*kn*_ is the *n*^*th*^ base or gap of *R*_*k*_ in the integrated MSA. The probability of a germline variant *p*(*S*_*kn*_|*R*_*kn*_) is calculated by

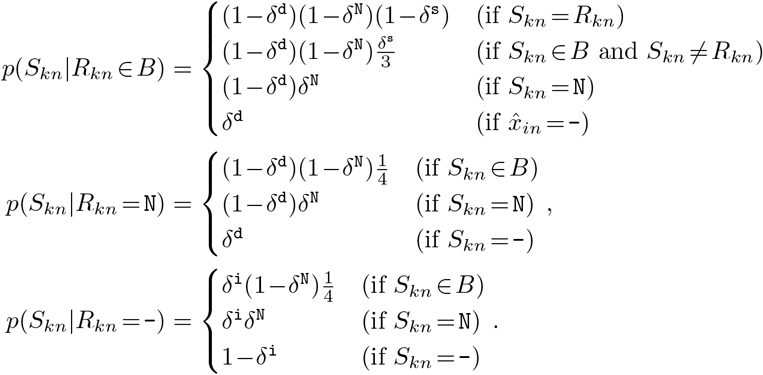

Here, δ^s^, δ^d^, δ^i^, and δ^N^ are the probabilities of a true substitution, a true deletion, a true insertion, and a true N, respectively. *S*_*kn*_ tends to become N when it is ambiguous.

The prior probability of the indicator variables 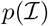 is defined by

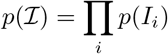

Here, *p*(*I*_*i*_) is the prior probability of the indicator variable, which is calculated by

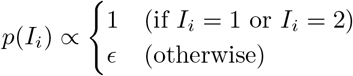

*ϵ* reflects how likely the reads are to be produced by non-target homologous regions.

### Efficient sampling with elaborate MCMC schemes

The parameters of the model above are sampled using two Markov chain Monte Carlo (MCMC) schemes, Gibbs sampling and the Metropolis-Hastings algorithm, with parallel tempering to make the parameter sampling efficient. Gibbs sampling is mainly used for local search, and Metropolis-Hastings sampling is periodically used for more global search. For the Metropolis-Hastings algorithm, we constructed two novel proposal distributions that enable the parameters to jump from mode to mode and lead more efficient sampling.

One of the proposal distributions is focused on positions not covered with any read. First, 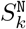 is defined as a modified HLA sequence whose bases are replaced with Ns at positions not covered with any read produced by *S*_*k*_, which is given by

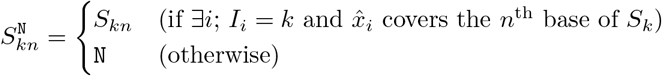

A candidate HLA type and a candidate HLA sequence are then sampled based on

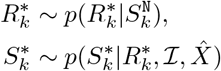

Then, the acceptance rate *r* can be calculated based on the Metropolis-Hastings algorithm, which is given by

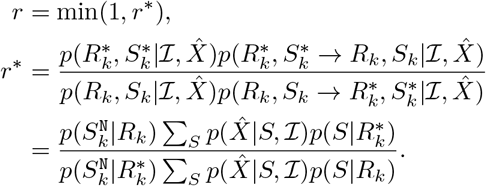

This proposal distribution makes the sampling more efficient when there is ambiguity in the HLA types attributed to some uncovered positions. For example, let *t*_*j*_ and *t*_*j*′_ be HLA types that only differ with one mismatch at the *n*^*th*^ position. If a sample has *t*_*j*_ as an HLA type but there are no reads from *t*_*j*_ covering the *n*^*th*^ position, we cannot determine whether the HLA type is *t*_*j*_ or *t*_*j*′_. However, once *R*_*k*_ becomes *t*_*j*′_, *S*_*kn*_ becomes the *n*^*th*^ base of *t*_*j*′_ with high probability. Then, *R*_*k*_ becomes *t*_*j*′_ with high probability, and this process is repeated. This is because *R*_*k*_ and *S*_*k*_ are separately sampled in the Gibbs sampling in spite of their high correlation. Thus, the proposal distribution prevents the parameters from getting stuck by sampling them simultaneously.

The other proposal distribution swaps non-decoy and decoy parameters. In this proposal distribution, indices for non-decoy and decoy parameters are uniformly sampled, and the HLA types and the HLA sequences at the indices are swapped. After swapping, candidate indicator variables are sampled based on the conditional distribution given the swapped parameters. Suppose that 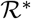 and 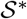 are HLA types and HLA sequences after swapping, and that 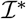 is a set of candidate indicator variables. Then, the acceptance rate *r* can be calculated by

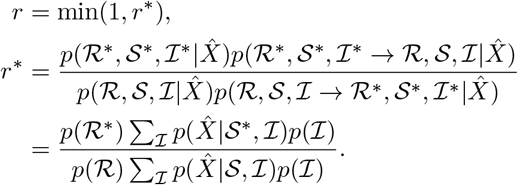

This proposal distribution enables quickly distinguishing reads from the target HLA gene and non-target homologous regions.

Some procedures are used in the burn-in period to avoid getting stuck in local optima. At the beginning of sampling, a multi-start strategy is used to reduce the influence of initial parameters. Specifically, some MCMC runs are carried out, and initial parameters are sampled from the last parameters of the MCMC runs. In addition, reference sequences are periodically copied to HLA sequences because there are many local optima where the parameters of the HLA sequences are twisted as if some crossovers occurred.

After sampling the parameters, HLA genotyping can be performed by counting 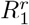 and 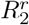. We used the most sampled HLA genotype in the MCMC process as the candidate. The HLA sequences of a sample can be also inferred by counting 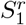 and 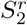.

## Results

### WES and WGS datasets

To evaluate the capability of our method, we obtained 253 WES data with the HLA genotypes from the International HapMap Project [22] that had been used by Szolek *et al.* [13] and Shukla *et al.* [15]. We further downsampled these data to 1/2, 1/4, 1/8, and 1/16 to simulate low-coverage data.

We also used paired normal and tumor WGS data of 25 Japanese cancer patients, including 20 liver cancer and 5 microsatellite-unstable colon cancer samples. These data were obtained from an Illumina HiSeq system with a 101-bp pair-end read length. The sequence data were deposited into the International Cancer Genome Consortium (ICGC) database (https://dcc.icgc.org/).

The sequencing-based typing (SBT) approach, which is guaranteed to be accurate at 4-digit resolution, was used for validation of the 20 liver cancer samples. Additional HLA genotyping using the TruSight HLA Sequencing Panels, which are theoretically guaranteed to be accurate at full (8-digit) resolution, was performed for 7 out of the above 20 liver cancer samples to reduce ambiguity of the SBT genotyping. The 5 microsatellite-unstable samples were genotyped using the TruSight HLA Sequencing Panels, in order to verify not only the HLA genotypes but also the presence of somatic mutations. We regarded the results of the SBT approach and/or the TruSight HLA Sequencing Panels as the correct information. If the results differed between the two methods, we assumed that the result of the TruSight HLA Sequencing Panel was correct.

### WES- and WGS-based HLA genotyping

For performance comparison, we used three existing methods, OptiType [13], PHLAT [12], and HLA-VBSeq [14] because it has been reported that they achieve the highest accuracy for WES- and WGS-based HLA genotyping [16]. First, we applied ALPHLARD and the existing methods to the original and the downsampled WES data (Additional file 1: Tables S1-S5). Because the gold standard HLA genotypes were determined from exon 2 and 3, we used only the exons as the reference sequences in ALPHLARD. Figure 2 shows the performance of the methods. ALPHLARD kept higher accuracy compared with the other methods even when the downsampling ratio was low. The accuracy of the existing methods was consistent with the preceding paper [16].

We also applied the methods to the normal WGS data and compared the determined HLA genotypes with those obtained by the SBT approach and the TruSight HLA Sequencing Panel (Additional file 2: Tables S6-S13). Table 1 shows the performance of the four methods. ALPHLARD clearly achieved a higher accuracy rate than the other methods. Moreover, the HLA-B genotype of one sample was inferred differently between the SBT approach and the TruSight HLA Sequencing Panel, and the result of ALPHLARD for this sample was identical to that of the TruSight HLA Sequencing Panel. This suggests that ALPHLARD could be potentially superior to the SBT approach in some cases. HLA-VBSeq achieved higher accuracy from the WGS data than from the WES data. This would be because HLA-VBSeq uses non-coding information such as the introns and the untranslated regions. The accuracy of the existing methods was consistent with the preceding paper [16].

**Table 1:** WGS-based HLA genotyping of ALPHLARD, OptiType, PHLAT, and HLA-VBSeq. N/A indicates that the method does not support the HLA gene.

|  |  | ALPHLARD | OptiType | PHLAT | HLA-VBSeq |
| --- | --- | --- | --- | --- | --- |
| HLA-A | 2-digit | **100% (50/50)** | **100% (50/50)** | 76.0% (38/50) | 96.0% (48/50) |
|  | 4-digit | **98.0% (49/50)** | **98.0% (49/50)** | 60.0% (30/50) | 82.0% (41/50) |
|  | 6-digit | **98.0% (49/50)** | N/A | 46.0% (23/50) | 82.0% (41/50) |
| HLA-B | 2-digit | **100% (48/48)** | 87.5% (42/48) | 72.9% (35/48) | 89.6% (43/48) |
|  | 4-digit | **100% (48/48)** | 85.4% (41/48) | 56.3% (27/48) | 75.0% (36/48) |
|  | 6-digit | **95.8% (46/48)** | N/A | 39.6% (19/48) | 72.9% (35/48) |
| HLA-C | 2-digit | **100% (50/50)** | **100% (50/50)** | 78.0% (39/50) | 96.0% (48/50) |
|  | 4-digit | **98.0% (49/50)** | 94.0% (47/50) | 56.0% (28/50) | 66.0% (33/50) |
|  | 6-digit | **98.0% (49/50)** | N/A | 44.0% (22/50) | 66.0% (33/50) |
| HLA-DPA1 | 2-digit | **100% (24/24)** | N/A | N/A | 87.5% (21/24) |
|  | 4-digit | **100% (24/24)** | N/A | N/A | 87.5% (21/24) |
|  | 6-digit | **100% (24/24)** | N/A | N/A | 87.5% (21/24) |
| HLA-DPB1 | 2-digit | **100% (22/22)** | N/A | N/A | 86.4% (19/22) |
|  | 4-digit | **100% (22/22)** | N/A | N/A | 86.4% (19/22) |
|  | 6-digit | **100% (22/22)** | N/A | N/A | 86.4% (19/22) |
| HLA-DQA1 | 2-digit | **100% (24/24)** | N/A | 70.8% (17/24) | **100% (24/24)** |
|  | 4-digit | **95.8% (23/24)** | N/A | 62.5% (15/24) | **95.8% (23/24)** |
|  | 6-digit | **95.8% (23/24)** | N/A | 62.5% (15/24) | **95.8% (23/24)** |
| HLA-DQB1 | 2-digit | **100% (18/18)** | N/A | 77.8% (14/18) | **100% (18/18)** |
|  | 4-digit | **94.4% (17/18)** | N/A | 61.1% (11/18) | 88.9% (16/18) |
|  | 6-digit | **94.4% (17/18)** | N/A | 38.9% (7/18) | 88.9% (16/18) |
| HLA-DRB1 | 2-digit | **100% (24/24)** | N/A | 70.8% (17/24) | 95.8% (23/24) |
|  | 4-digit | **100% (24/24)** | N/A | 50.0% (12/24) | 58.3% (14/24) |
|  | 6-digit | **100% (24/24)** | N/A | 45.8% (11/24) | 58.3% (14/24) |
| Total | 2-digit | **100% (260/260)** | 95.9% (142/148) | 74.8% (160/214) | 93.8% (244/260) |
|  | 4-digit | **98.5% (256/260)** | 92.6% (137/148) | 57.5% (123/214) | 78.1% (203/260) |
|  | 6-digit | **97.7% (254/260)** | N/A | 45.3% (97/214) | 77.7% (202/260) |

**Figure 2:**
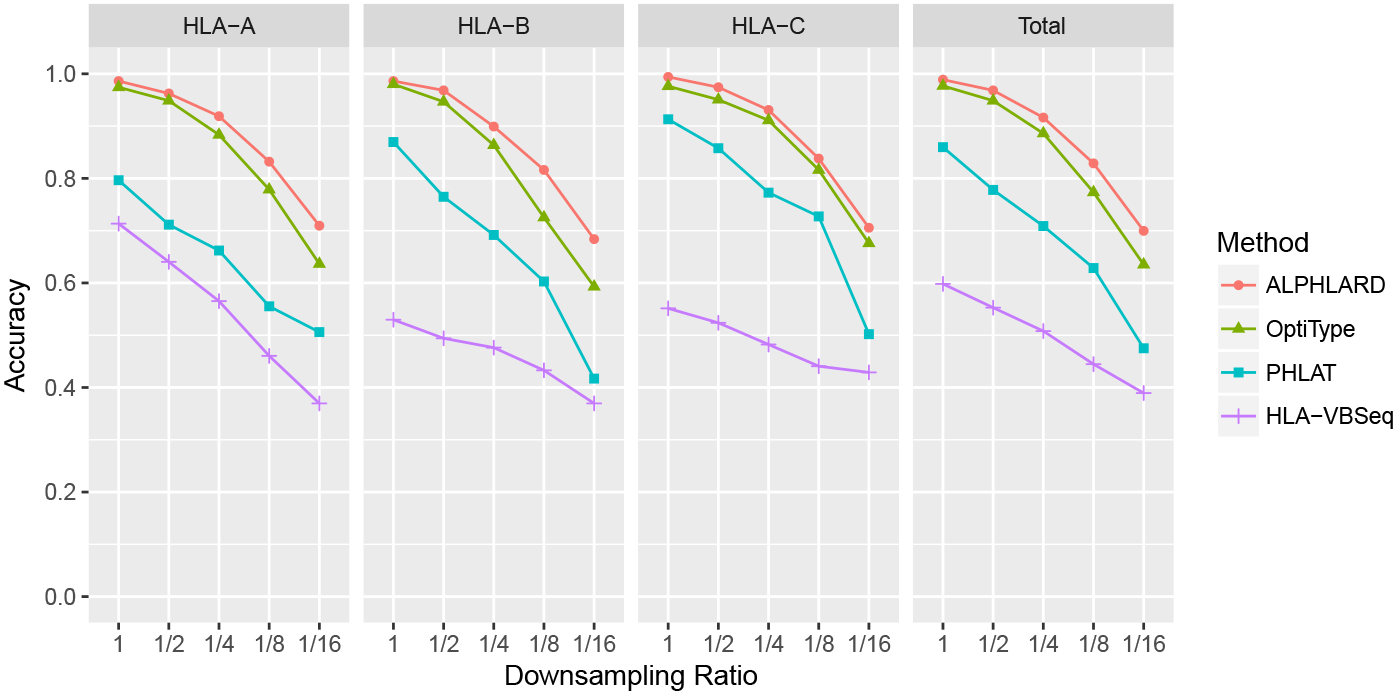
WES-based HLA genotyping of ALPHLARD, OptiType, PHLAT, and HLA-VBSeq. Each WES data was downsampled to 1/2, 1/4, 1/8, and 1/16, and the four methods were applied to all of the original and the downsampled WES data.

### Detection of somatic mutations

Next, we searched for somatic point mutations in the HLA genes. They were detected by comparing the inferred HLA sequences between paired normal and tumor samples of each patient. We detected three somatic point mutations in the microsatellite-unstable samples: two single-base deletions and one singlebase insertion (Figure 3 and Additional file 3: Figures S1 and S2). One of the deletions occurred in a homopolymeric region in exon 1 of the HLA-A gene, and the other occurred in a homopolymeric region in exon 1 of the HLA-B gene. Both of these mutations caused a frameshift, leading to an early stop codon and ultimate loss of function of the HLA allele. It is known that the HLA-A and HLA-B genes are homologous, and we found that the two deletions occurred at homologously the same position. Moreover, one of the HLA-A types (A*68:11N) has a single-base deletion at exactly the same homopolymeric position. These observations suggest that the homopolymeric regions are deletion hotspots. The insertion occurred in a homopolymeric region at the beginning of exon 4 of the HLA-A gene, which changed the HLA-A allele from A*31:01:02 to A*31:14N. This region is known as an insertion hotspot in some HLA types such as A*01:04N and B*51:11N, and the insertion causes no expression of the allele [23–26]. The three indels identified were validated by the TruSight HLA Sequencing Panels and the Sanger sequencing.

**Figure 3:**
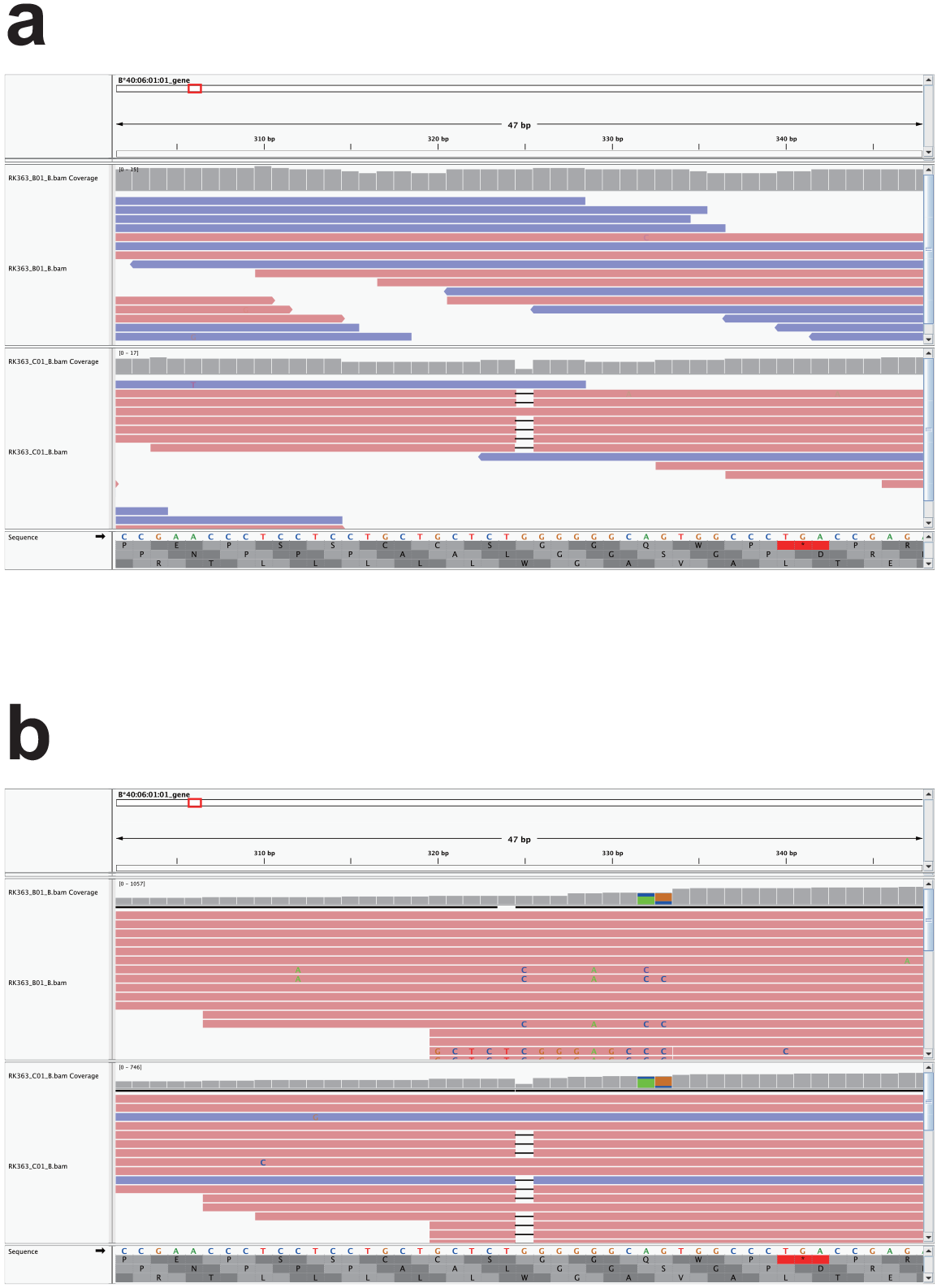
A single-base deletion in exon 1 of the HLA-B gene of patient RK363. IGV screenshots were taken at the position for (a) the WGS data and (b) the TruSight HLA Sequencing Panel data. In each of the screenshots, the upper and lower tracks correspond to the normal and tumor samples, respectively.

We further sought cases of LOH in the HLA genes as follows. First, we focused on two types of patients: (i) those for which HLA genotypes were uniquely determined for the normal sample but not for the tumor sample, and (ii) those for which HLA genotypes of both the normal and the tumor samples were uniquely but not identically determined. Then, we checked whether the collected reads of the tumor sample supported the HLA genotype inferred for the normal sample.

We were able to detect one likely case of LOH in the tumor sample of a patient, RK069. At each heterozygous single nucleotide polymorphism (SNP) position in each HLA locus, the log odds ratio was calculated for the WGS data and the Trusight HLA Sequencing Panels based on the number of reads that supported the SNP (Figure 4 and Additional file 4: Figures S3-S7). These figures suggest that A*26:01:01, B*35:01:01, C*03:03:01, DPA1*01:03:01, DQA1*03:02, and DRB1*12:01:01 might be lost in the tumor sample of RK069.

## Discussion

In this paper, we presented a new Bayesian method, ALPHLARD, which performs not only HLA genotyping but also infer the HLA sequences of a sample. The results showed that our method ALPHLARD achieved higher accuracy for HLA genotyping from both WES and WGS data than existing methods. We presume that the high performance of ALPHLARD originates from the following reasons. First, the search space of ALPHLARD is all possible HLA allele pairs. Some methods treat an HLA allele pair as two independent HLA alleles; that is they give a score to each HLA allele and output the most and the second most probable HLA alleles without directly considering the combinations. This approximation reduces the computation time but works well only when the coverage of the sequence data is sufficient. Therefore, such methods would not achieve high accuracy for HLA genotyping from WGS data. Second, ALPHLARD takes into account whether or not bases and gaps are observed at each position by inserting the parameters for HLA sequences between the parameters for HLA genotypes and collected reads. Most of read count-based HLA genotyping algorithms consider only the number of reads mapped to each HLA allele. However, even if a lot of reads are mapped to an HLA allele, it does not seem to be the true HLA type if there are several regions not covered by any read. We believe that what is really important is not the number of reads but the range covered by sufficient reads. Third, ALPHLARD uses some decoy parameters in addition to non-decoy ones. This is why ALPHLARD can robustly and accurately perform HLA genotyping even if there exist some reads from non-target homologous regions that are similar to the target HLA gene.

Besides HLA genotypes, ALPHLARD gives us beneficial information that cannot be obtained from other methods. First, somatic mutations such as point mutations and LOHs can be detected by comparing the sampled HLA sequences of paired normal and tumor samples. We detected three indels and one case of LOH, which lead to loss of function of the HLA alleles. These mutations are biologically important because they weaken the immune function and would be related to tumor progression. Second, novel HLA types not registered in HLA databases can be identified by comparing the inferred HLA genotype and HLA sequences. Unfortunately, no novel HLA type was observed in our analysis. However, ALPHLARD would be flexible enough to detect the difference between novel HLA types and known ones because the process of novel HLA type identification is theoretically the same as that of HLA somatic mutation detection.

**Figure 4:**
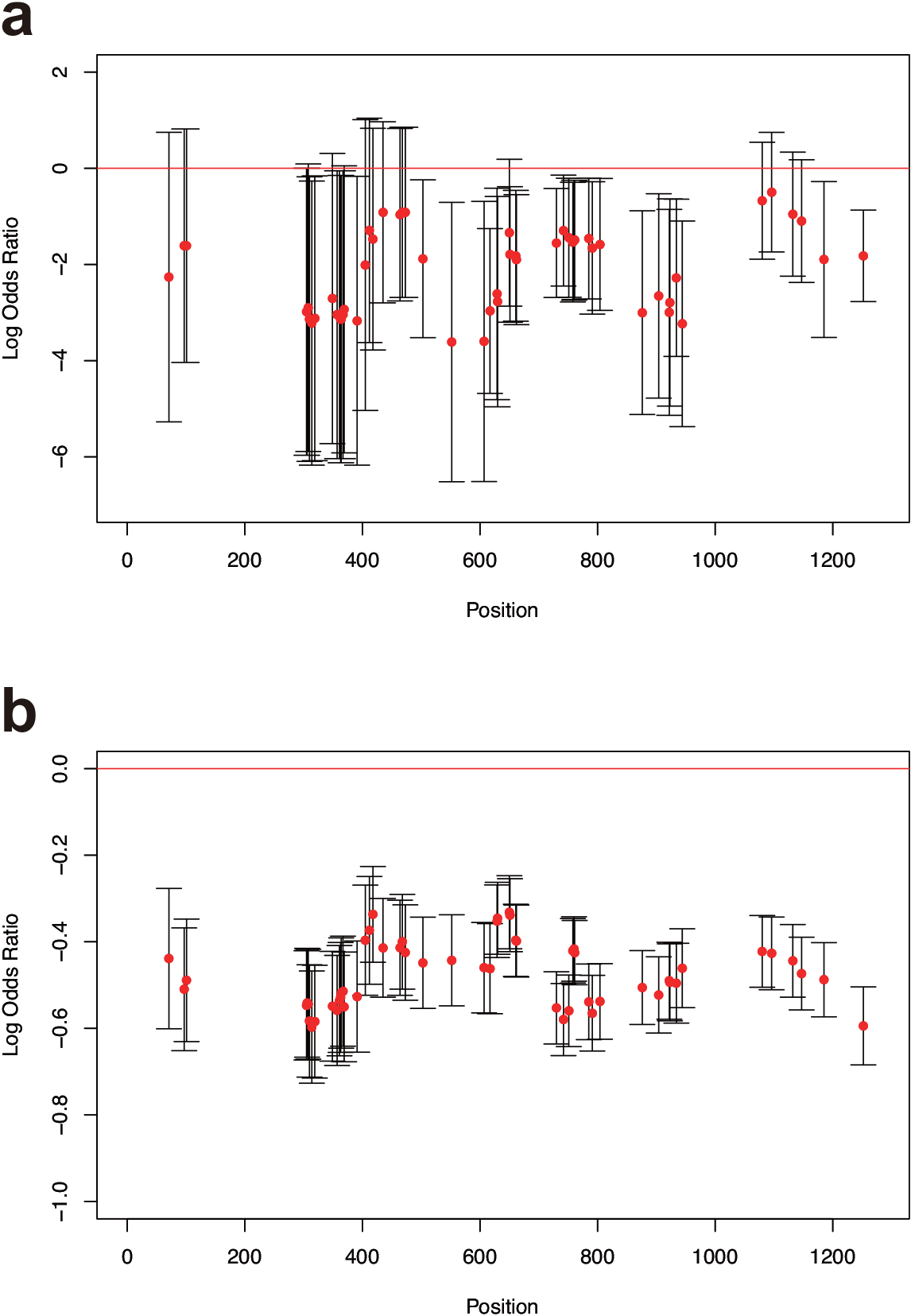
The log odds ratios of the depths at heterozygous SNP positions in the HLA-A gene of patient RK069. The log odds ratios were calculated for (a) the WGS data and (b) the TruSight HLA Sequencing Panel data. These log odds ratios correspond to the relative quantities of observed A*26:01:01 SNPs in the tumor sample compared with the normal sample. The red dots indicate the mean values of the log odds ratios, and the vertical lines indicate the 95% confidence intervals.

## Conclusion

Our new Bayesian-based HLA analysis method, ALPHLARD, showed good performance for HLA genotyping. It also has a potential to detect rare germline variants and somatic mutations in HLA genes. A large amount of WGS data has been recently produced by big projects such as the ICGC. Applying our method to such big data would help to fill in the current gaps in HLA reference databases and unveil the immunological significance of somatic mutations identified in HLA genes.

## Abbreviations

HLA: human leukocyte antigen; LOH: loss of heterozygosity; MCMC: Markov chain Monte Carlo; MSA: multiple sequence alignment; SBT: sequencing-based typing; SNP: single nucleotide polymorphism; WES: whole exome sequencing; WGS: whole genome sequencing;

## Ethics approval and consent to participate

All of the human subjects agreed with informed consent to participate in the study following ICGC guidelines [27]. IRBs at RIKEN and the associated hospitals participating in this study approved this work.

## Consent for publication

Not applicable.

## Availability of data and material

The WGS data were deposited into the ICGC database (https://dcc.icgc.org/).

## Competing interests

The authors declare that they have no competing interests.

## Funding

This work was supported by Japan Society for the Promotion of Science (15H02775 and 15H05912).

## Authors’ contributions

SH, RY, SM, and SI designed the research. SH developed the method. SH and MK benchmarked the method. MS performed SBT genotyping to the samples. HN provided the whole genome and the amplicon sequencing data of the samples. SH and SI wrote the manuscript. All authors read and approved the final manuscript.

## Acknowledgements

The super-computing resource was provided by Human Genome Center, the Institute of Medical Science, the University of Tokyo.

